# A chromosome-level reference genome of the largest cervid species - the European moose (*Alces alces*; Linnaeus, 1758)

**DOI:** 10.64898/2026.07.03.736352

**Authors:** Ole K. Tørresen, Atle Mysterud, Morten Skage, Bram Danneels, Marius A. Strand, Giada Ferrari, Ave Tooming-Klunderud, Kjetill S. Jakobsen

**Affiliations:** Dept of Biosciences, Centre for Ecological and Evolutionary Synthesis (CEES), University of Oslo, Oslo, Norway; Computational Biology Unit, Department of Informatics, University of Bergen, Bergen 5020, Norway

## Abstract

We describe a chromosome-level, haplotype-resolved genome assembly from a male European moose (*Alces alces alces*). The assembly comprises two pseudo-haplotypes of 3,148 Mb and 3,112 Mb, with sex chromosomes in haplotype one, and 33 autosomes in each haplotype (68 in total). Assembly completeness is high (BUSCO 98.3% and 95.7%), with 21,496 and 20,498 annotated protein-coding genes for haplotypes one and two, respectively. This genome assembly is the most complete so far generated for European moose.

## Introduction

The moose (*Alces alces;* Linnaeus, 1758), termed elk in British English, is the largest extant member of the family Cervidae and an iconic species across the Northern Hemisphere. Its circumpolar distribution cover about 26.2 million km^2^ spanning the boreal forest biome including taiga and low-elevation mountains of Eurasia from Scandinavia and Poland eastward through Siberia to northern China and Mongolia (Jensen et al., 2020; Rodgers et al., 2025). Moose immigrated to North America from Asia through Beringia only some 14,000-11,000 years ago (Hundertmark et al., 2002). Today in North America, moose are present from Alaska and Canada southward along northern parts of the Rocky Mountains in the USA, and eastwards in Canada and northern USA towards the Atlantic Ocean (A. W. Franzmann & Schwartz, 1997; Rodgers et al., 2025). The postharvest population in North America is estimated above 1,100,000 individuals, while it is around 1,200,000 moose in E. The species is cold-adapted (Niedziałkowska et al., 2024; Schwab & Pitt, 1991; Telfer & Kelsall, 1984) and struggle with effects of climate change in the southern parts of its distribution range both in North America (Lenarz et al., 2009; Murray et al., 2006) and Europe (Holmes et al., 2021).

Moose have a high stature with long legs and body masses often reaching 300-500 kg among adults, and they can occasionally reach above 700 kg (Rodgers et al., 2025). Moose is weakly polygynous with moderate sexual body size dimorphism (Loison et al., 1999; Markussen et al., 2019). Prime-aged moose cows regularly give birth to twins under good conditions (Albert W. Franzmann & Schwartz, 1985), which is uncommon for a large herbivore (Gaillard, 2007). Moose is a typical browser, favoring early successional forest stages and riparian zones rich in browse species such as willow (*Salix* spp.), birch (*Betula* spp.), Scots pine (*Pinus sylvestris*), and *Vaccinium* spp. (Shipley et al., 1998; Spitzer et al., 2023). During summer, diet also includes a range of forbs as well as aquatic plants especially in North America (Belovsky, 1981). The availability of high-quality browse largely determines the body condition and thereby reproduction. Moose are considered a keystone herbivore shaping forest composition and influencing nutrient cycling through selective feeding (Danell et al., 1994; Pastor & Naiman, 1992). Furthermore, moose have been integral to human subsistence and culture (Rosvold et al., 2013), and had a wider distribution in Europe previous times (Schmölcke & Zachos, 2005). Archaeological evidence and historical records indicate that moose hunting has provided meat, hides, and antlers for thousands of years (Hall, 1973; Regelin & Franzmann, 1998), and the species remains an important game animal in many regions today (Lavsund et al., 2003; Natcher et al., 2021).

Although taxonomic debates persist (Hundertmark, 2016), *Alces alces* is generally treated as a single species. The number of subspecies is more debated (Rodgers et al., 2025), or whether variations only reflect ecotypes. Some authors describe up to eight subspecies with four on each continent . European and West Siberian moose possess 68 chromosomes, while East Asian and North American moose have 70 chromosomes (Boeskorov, 1997). Current evidence from marking studies points to a high rate (10-30%) of juvenile dispersal in both sexes and often long dispersal distances (Ballard et al., 1991; Cederlund et al., 1987; Dodge et al., 2004; Labonte et al., 1998). Consequently, there is limited spatial genetic structure at local scales both in North America (Rosenblatt et al., 2023; Schmidt et al., 2009) and Europe (Niedziałkowska et al., 2016). In North America, there was genetic differentiation between some regions linked to major barriers like large rivers and mountain ranges in North America (Hundertmark et al., 2003; Rosenblatt et al., 2023) and similarly suggested by subspecies listing for Eurasia (Baskin & Danell, 2003). Across Europe, the two main genetic clusters are continental versus Scandinavian moose, reflecting a split during the last glaciation and with the Baltic Sea as a barrier to gene flow. There is a differentiation between these European populations towards Russia and Asia (Świsłocka et al., 2020), but detailed studies are lacking for most of Asia and genetic borders towards Europe are not distinct. In Scandinavia, there is a further split between and southern parts of both Sweden (Dussex et al., 2020, 2023; Wennerström et al., 2016) and Norway (Haanes et al., 2011) with an intermediate mixing zone.

Here, we present a chromosome-level genome assembly of European moose sampled in southeast Norway and generated by long read sequencing (PacBio HiFi) and scaffolding by Hi-C. This genome represents a valuable resource for population genomic studies, and studies addressing the genetics of climate effects as well as comparative studies of cervids. This genome assembly was generated as part of the Earth BioGenome Project Norway.

## Material and Methods

### Sample acquisition and DNA extraction

A heart tissue sample from a young male bull with spike antlers (*Alces alces)* was collected after being legally shot 9th of October 2023 during the annual hunting season at Aslum, Nordre Follo, Norway (sampling coordinates: 59.6421132, 10.8984526). During the butchering process the heart tissue sample was cut into smaller pieces and stored in 96% ethanol before transfer to the laboratory for long term storage at -80°C. Isolate name is EBPN_MS_8038_01 with specimen name AlcAlc071023 under the Earth Biogenome Norway initiative ([EBP-Nor](https://www.ebpnor.org/)).

The DNA isolation for PacBio long-read sequencing started from 109 mg ethanol preserved tissue which were taken from cold storage, blotted dry on tissue paper and transferred to CryoPrep (Covaris Inc) sample bags before cooling in liquid nitrogen and crushed with moderate force using three impacts on setting 3 on the Cryoprep. DNA was isolated from the crushed tissue using two 100G columns and the Genomic Tip kit from Qiagen (Qiagen N.V.). Quality check of the amount, purity and integrity of the isolated DNA was performed using a combination of Qubit BR DNA quantification assay kit (Thermo Fisher), Nanodrop (Thermo Fisher), and Fragment Analyser (DNA HS 50kb large fragment kit, Agilent Tech.).

### Library preparation and sequencing for *de novo* assembly

Before PacBio HiFi library preparation, DNA was purified an additional time using AMPure PB beads (1:1 ratio). Purified HMW DNA was sheared two times to a modal fragment size of 21 kbp large fragments using the Megaruptor3 (Diagenode) on speedcode setting 30+31 . A HiFi library was prepared following the PacBio protocol for HiFi library preparation using the SMRTbell^®^ Prep Kit 3.0. The final HiFi library was size-selected with a 10 kbp cut-off using a BluePippin (Sage Sciences) and sequenced on one SMRT cell (using the associated Revio Binding kit and Sequencing chemistry) on the PacBio Revio instrument (PacBio Inc) located at the Norwegian Sequencing Centre (University of Oslo, Norway).

For scaffolding of long reads we prepared a Hi-C library starting with 103 mg ethanol preserved tissue into the Arima High Coverage HiC kit (Arima Genomics), with the manufacturer’s user guide for standard input preparations (Document Part Number A160162 v01), followed by Illumina short read library preparation with the Arima library prep Module and associated user guide (A101030, A303011, A160186 v02). Final library quality was validated using the equipment mentioned above and quantified using a Kapa Library quantification kit for Illumina (Roche Diagnostics) on a LightCyler-96 qPCR instrument (Roche Diagnostics) . The library was sequenced with other libraries on an Illumina NovaSeq 6000 (lllumina Inc) with 2*150 bp paired end mode at the Norwegian Sequencing Centre (https://www.sequencing.uio.no).

### Genome assembly and curation, annotation and evaluation

A full list of relevant software tools and versions is presented in Table 1. We assembled the species using a pre-release of the EBP-Nor genome assembly pipeline (https://github.com/ebp-nor/GenomeAssembly). KMC (Kokot et al., 2017) was used to count k-mers of size 32 in the PacBio HiFi reads, excluding k-mers occurring more than 10,000 times. GenomeScope (Ranallo-Benavidez et al., 2020) was run as part of the pipeline on the k-mer histogram output from KMC and was included in the methods for completeness. Ploidy level was investigated using Smudgeplot . HiFiAdapterFilt (Sim et al., 2022) was applied on the HiFi reads to remove possible remnant PacBio adapter sequences. The filtered HiFi reads were assembled using hifiasm (Cheng et al., 2021) with Hi-C integration resulting in a pair of haplotype-resolved assemblies, pseudo-haplotype one (hap1) and pseudo-haplotype two (hap2). Unique k-mers in each assembly/pseudo-haplotype were identified using meryl (Rhie et al., 2020) and used to create two sets of Hi-C reads, one without any k-mers occurring uniquely in hap1 and the other without k-mers occurring uniquely in hap2.

**Table 1.**
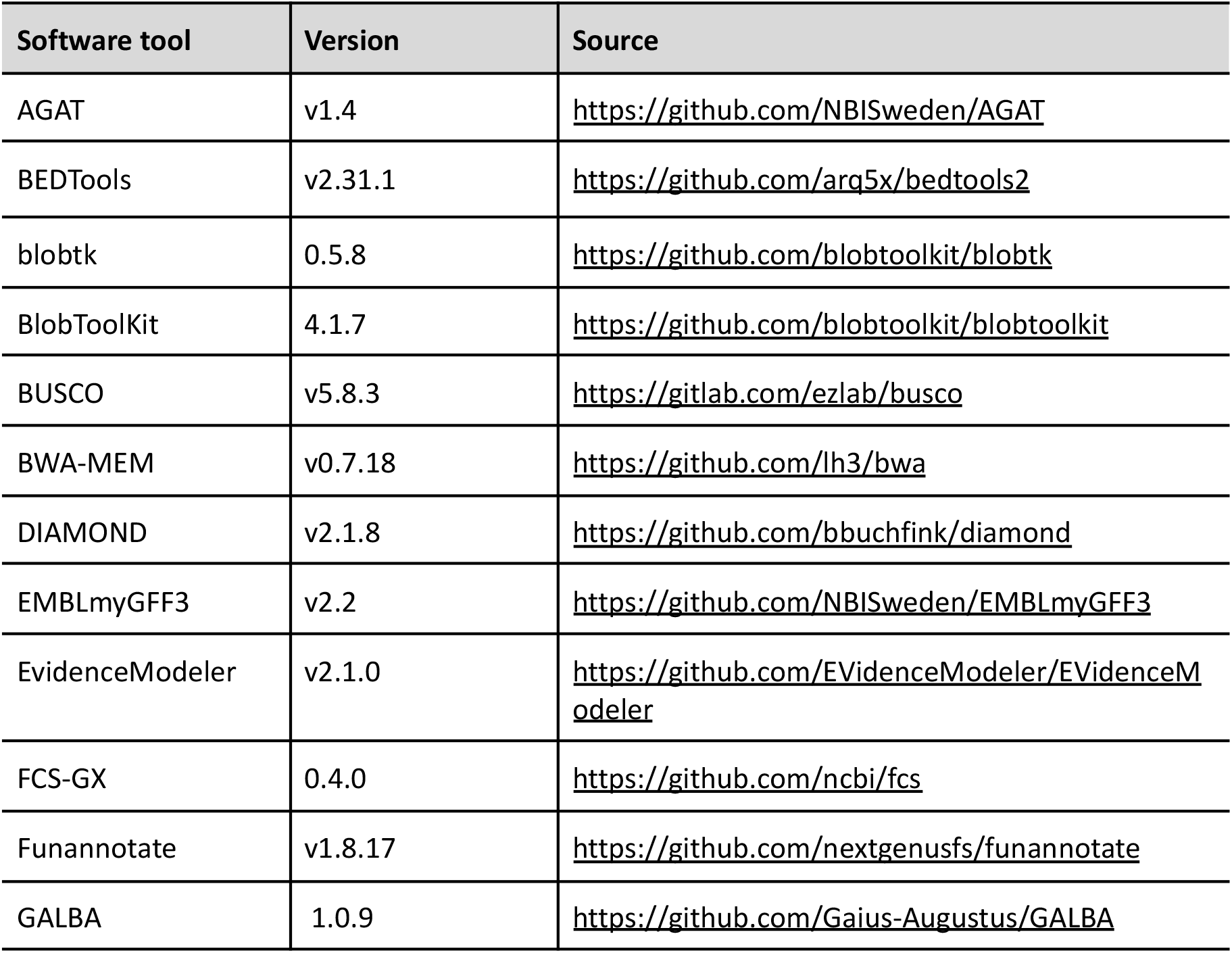
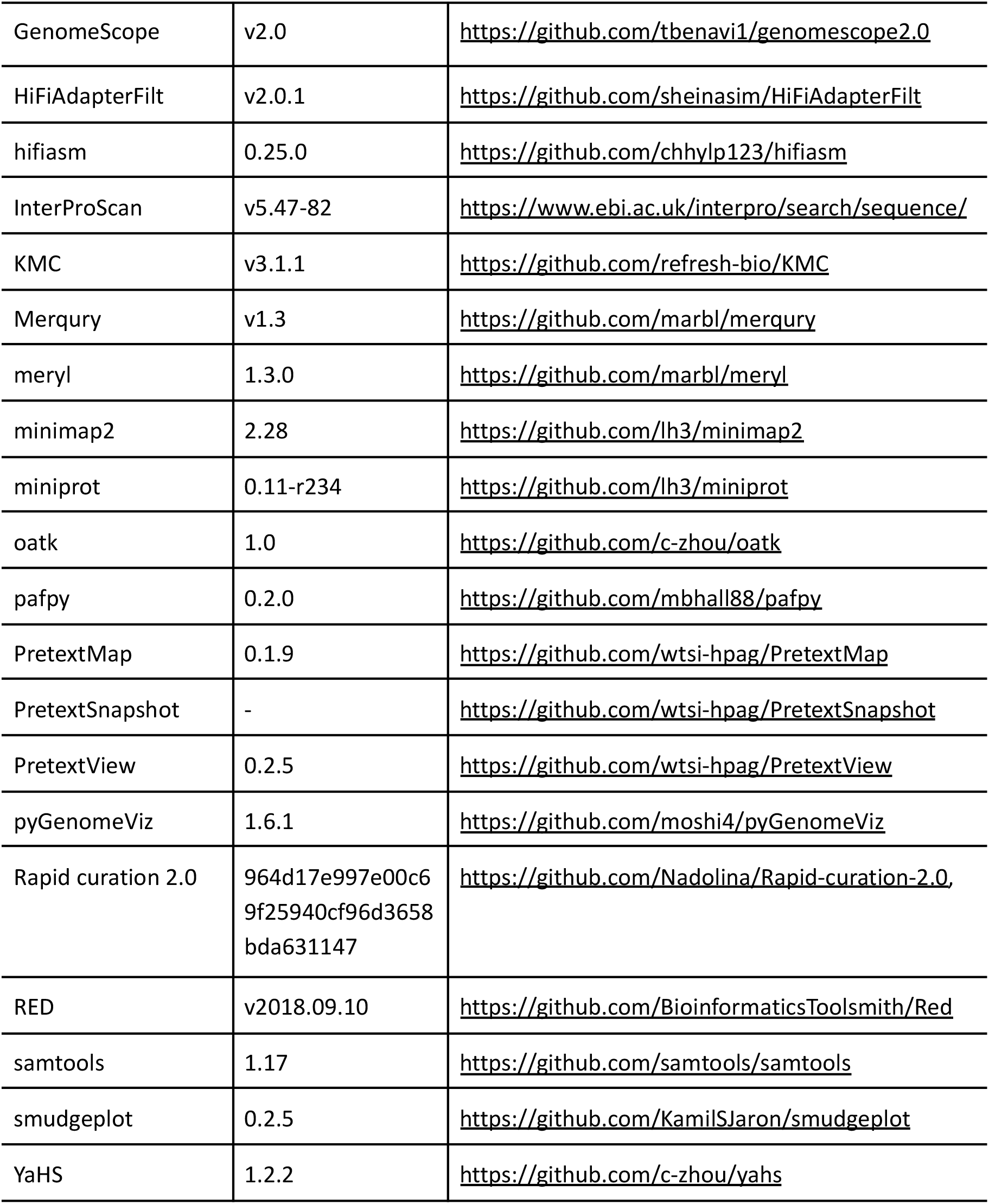
Software tools: versions and sources.

K-mer filtered Hi-C reads were aligned to each scaffolded assembly using BWA-MEM (Li, 2013) with -5SPM options. The alignments were sorted based on name using samtools (Li et al., 2009) before applying samtools fixmate to remove unmapped reads and secondary alignments and to add mate score, and samtools markdup to remove duplicates. The resulting BAM files were used to scaffold the two assemblies using YaHS (Zhou et al., 2023) with default options. FCS-GX (Astashyn et al., 2024) was used to search for putative contamination and contaminated sequences were removed. In addition, contigs with a HiFi coverage <7.5x or shorter than 1 Mb were removed from the assembly. The mitochondrion was searched for in reads using Oatk (Zhou et al., 2024). The assemblies were manually curated using PretextView and Rapid curation 2.0. Chromosomes were identified by inspecting the Hi-C contact map in PretextView. The sex chromosomes and the mitochondrial genome was added to haplotype one.

We annotated the genome assemblies using a pre-release version of the EBP-Nor genome annotation pipeline (https://github.com/ebp-nor/GenomeAnnotation). First, AGAT (https://zenodo.org/record/7255559) agat_sp_keep_longest_isoform.pl and agat_sp_extract_sequences.pl were used on the *Homo sapiens* (GRCh38) genome assembly and annotation to generate one protein (the longest isoform) per gene. Miniprot (Li, 2023) was used to align the proteins to the curated assemblies. UniProtKB/Swiss-Prot (UniProt Consortium, 2023) release 2025_3 in addition to the Mammalia part of OrthoDB v12 (Kuznetsov et al., 2023) were also aligned separately to the assemblies. RED (Girgis, 2015) was run via redmask (https://github.com/nextgenusfs/redmask) on the assemblies to mask repetitive areas. In addition to gene models from protein alignments, GALBA (Brůna et al., 2023; Buchfink et al., 2015; Hoff & Stanke, 2019; Li, 2023; Stanke et al., 2006) was run with the *Homo sapiens* proteins using the miniprot mode on the masked assemblies to generate *ab initio* predicted gene models. In addition, Helixer (Holst et al., 2025) was also run using the vertebrate-specific model (vertebrate_v0.3_m_0080).

The funannotate-runEVM.py script from Funannotate was used to run EvidenceModeler (Haas et al., 2008) on the alignments of GRCh38 proteins, UniProtKB/Swiss-Prot proteins, Mammalia proteins and the predicted genes from GALBA and Helixer. The resulting predicted proteins were compared to the protein repeats that Funannotate distributes using DIAMOND blastp and the predicted genes were filtered based on this comparison using AGAT. The filtered proteins were compared to the UniProtKB/Swiss-Prot release 2025_03 using DIAMOND (Buchfink et al., 2015) blastp to find gene names and InterProScan (Jones et al., 2014) was used to discover functional domains. AGATs agat_sp_manage_functional_annotation.pl was used to attach the gene names and functional annotations to the predicted genes. EMBLmyGFF3 (Norling et al., 2018) was used to combine the fasta files and GFF3 files into a EMBL format for submission to ENA.

All the evaluation tools have also been implemented in a pipeline, similar to assembly and annotation (https://github.com/ebp-nor/GenomeEvaluation). Merqury (Rhie et al., 2020) was used to assess the completeness and quality of the genome assemblies by comparing them to the k-mer content of both the Hi-C reads and PacBio HiFi reads. BUSCO (Manni et al., 2021) was used to assess the completeness of the genome assemblies by comparing against the expected gene content in the eudicots lineage. Gfastats (Formenti et al., 2022) was used to output different assembly statistics of the assemblies.

BlobToolKit and BlobTools2 (Laetsch & Blaxter, 2017), in addition to blobtk were used to visualize assembly statistics. To generate the Hi-C contact map image, the Hi-C reads were mapped to the assemblies using BWA-MEM (Li, 2013) using the same approach as above. Finally, PretextMap was used to create a contact map which was visualized using PretextSnapshot.

To compare the assemblies generated in this study to other available European moose genome assemblies, we downloaded GCA_964270355.1, GCA_015832495.2 and GCA_007570765.1 from NCBI and aligned these to each other and the assemblies generated here using minimap2 (Li, 2017) resulting in pairwise PAF files. For visualisation purposes, scaffolds in the downloaded assemblies were ordered according to hap1, and only scaffolds larger than 5 Mb were included in plots, or only chromosomes if that information was available. PAF files were processed using pafpy and pyGenomeViz was used to plot (see Zenodo repository listed under Data availability for the script used to plot).

Parts of the text in Methods and Results is based on a template we use for all the species we publish in the EBP-Nor project.

## Results

### *De novo* genome assembly and annotation

The genome of a male moose was assembled from a total of 23-fold coverage in Pacific Biosciences single-molecule HiFi long reads and 78-fold coverage in Arima Hi-C reads resulting in two haplotype-separated assemblies. The final assemblies have total lengths of 3,433 Mb and 3,113 Mb (Table 2 and Figure 2), respectively. Pseudo-haplotypes one (hap1) and two (hap2) have scaffold N50 size of 66.8 Mb and 60.8 Mb, respectively, and contig N50 of 28.6 Mb and 23.4 Mb, respectively (Table 2 and Fig. 2). 33 pseudo-chromosomes were identified in both pseudo-haplotypes, in addition to sex chromosomes (X and Y) in hap1. Chromosomes are named according to size and the Hi-C contact maps show clear separation of chromosomes into homologous sets (Supplementary Figure 1).

**Table 2:**
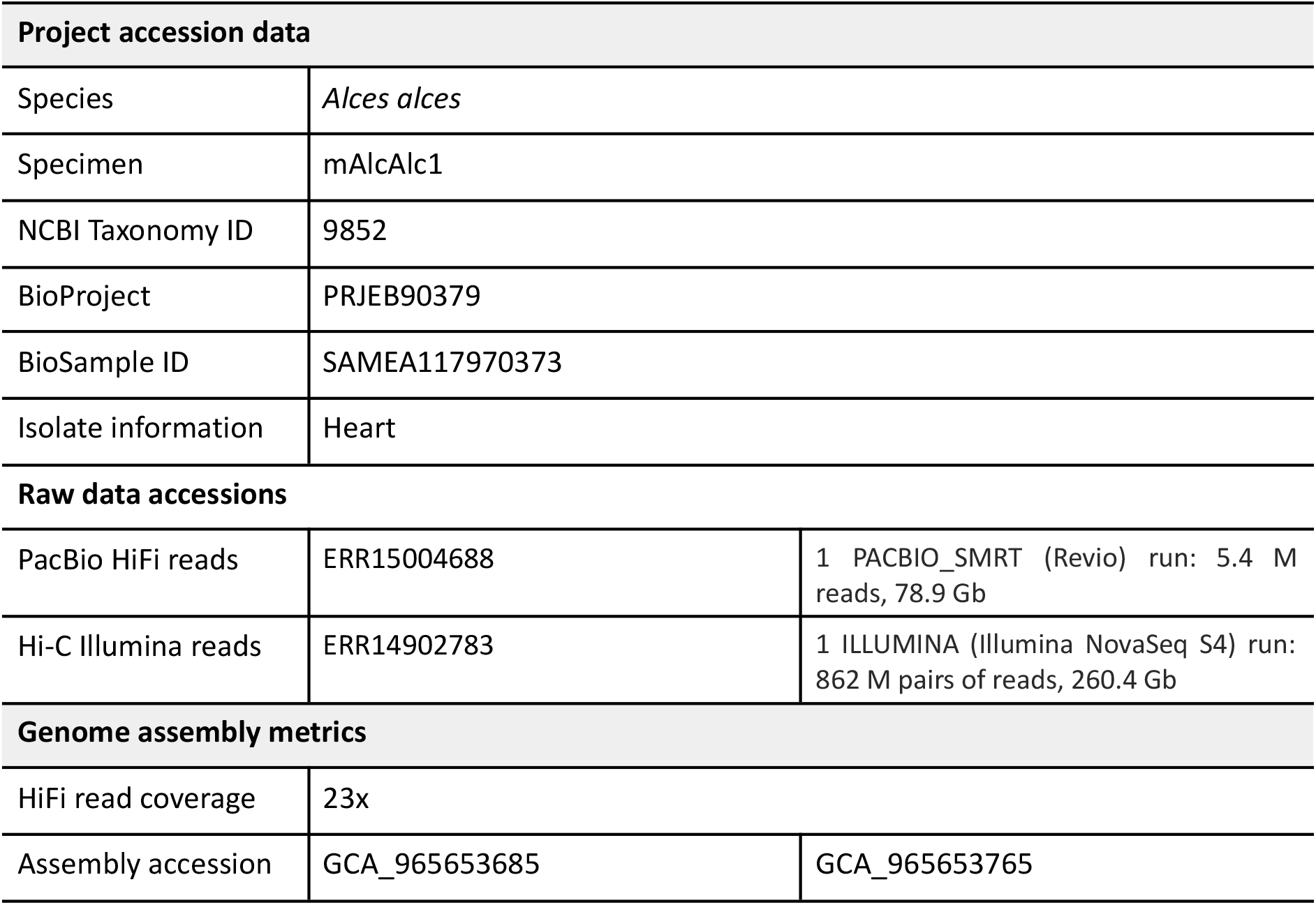
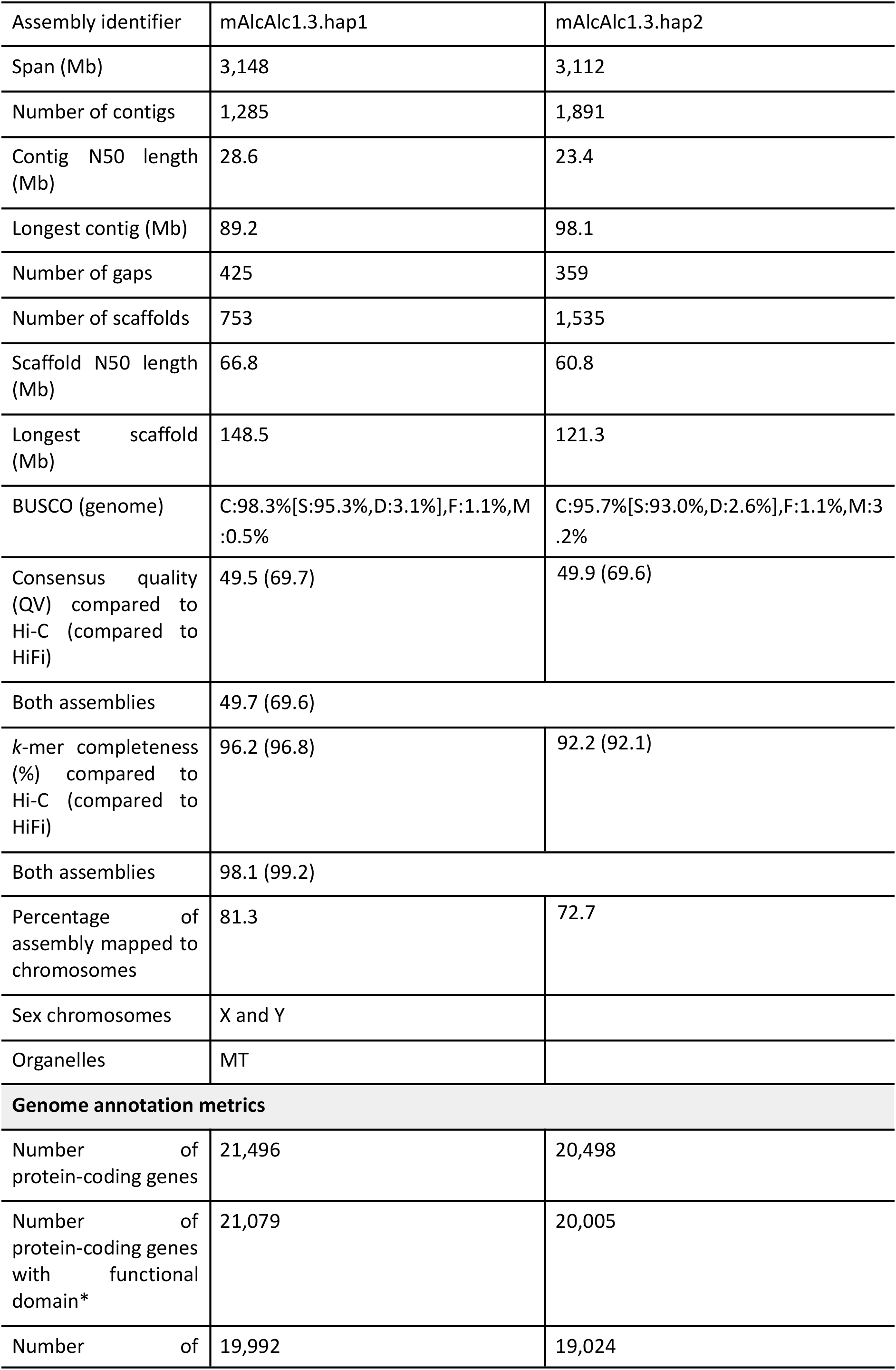
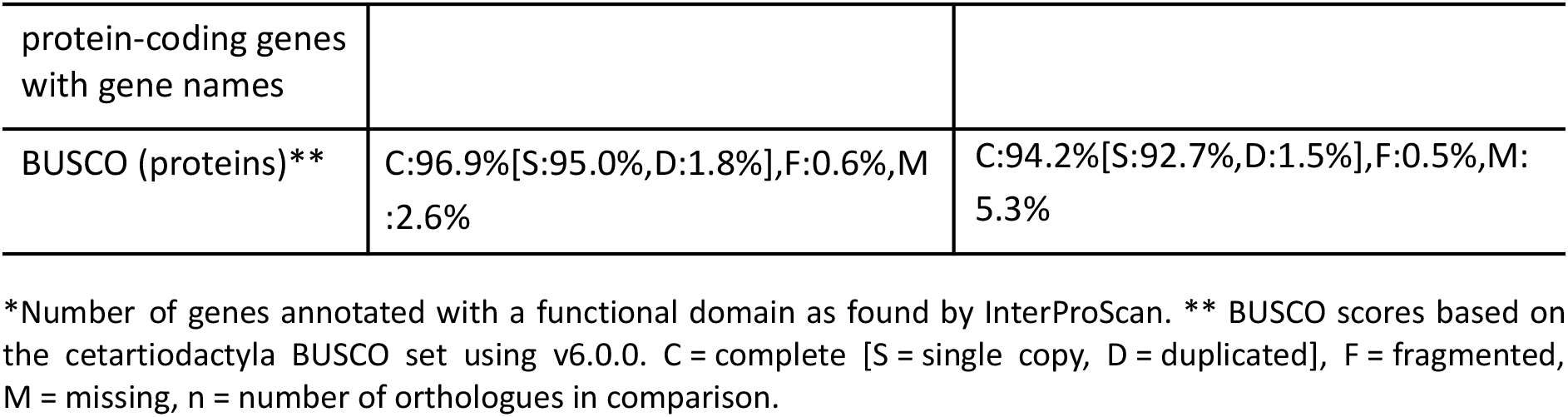
Genome data for *Alces alces*.

**Figure 1.**
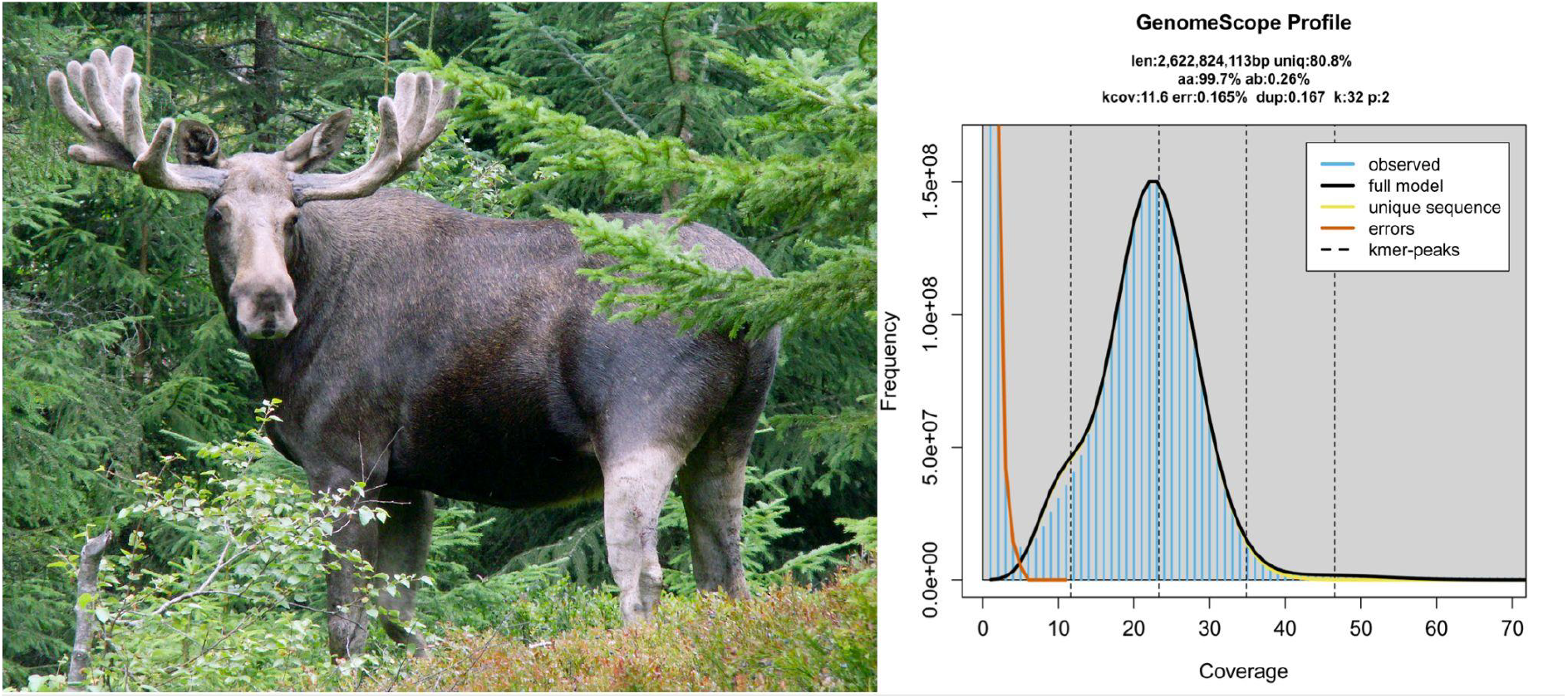
Example specimen and genome profile. Picture of adult male European moose (by Badzil, licensed under CA BY-SA 3.0) and GenomeScope profile of the HiFi reads from the sequenced individual. This analysis estimates a 2,622 Mb genome, with 0.26% heterozygosity. The left-hand peak of k-mers corresponds to k-mers from heterozygous regions of the genome, while the right-hand peak is from homozygous regions.

**Figure 2:**
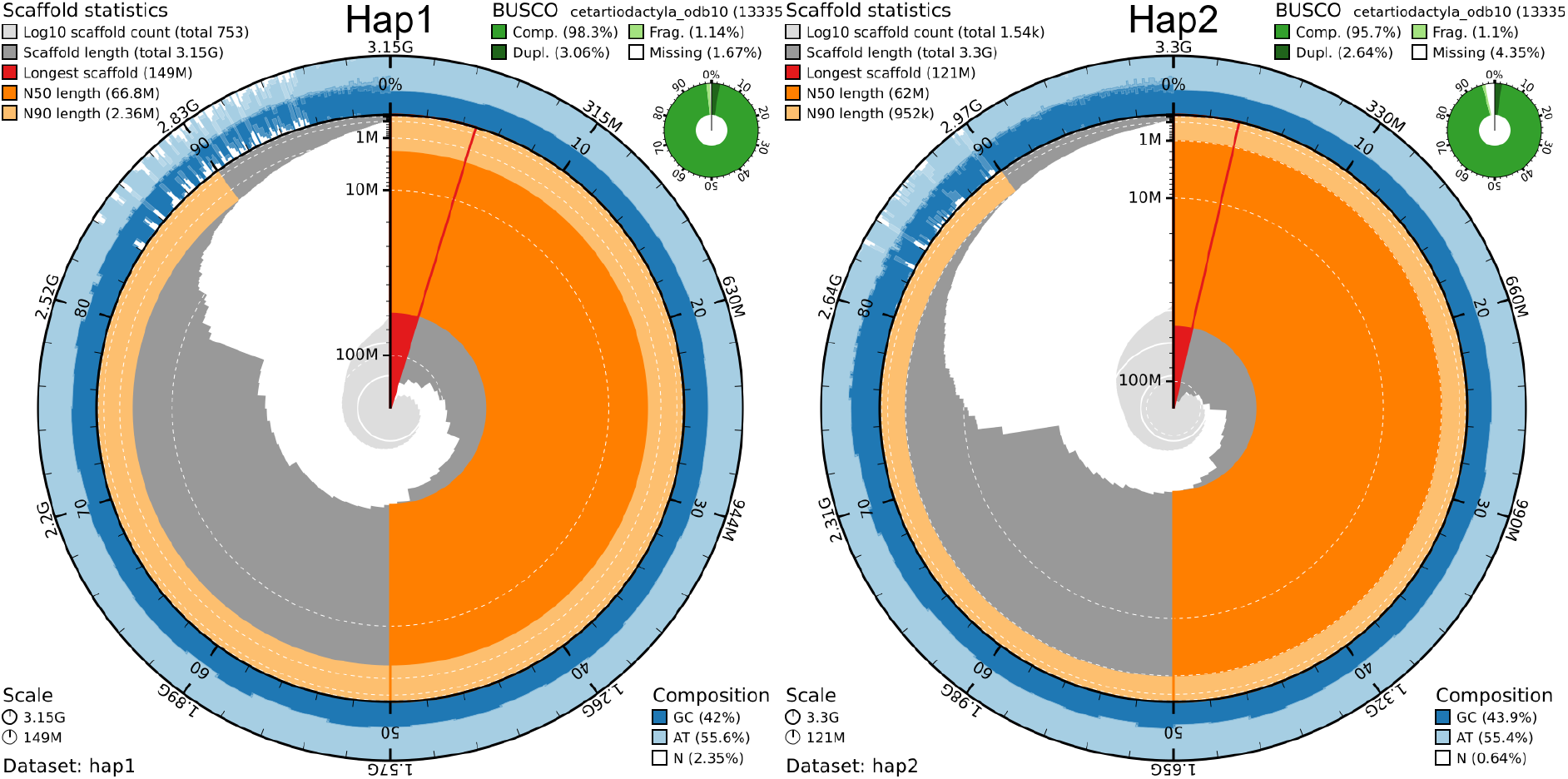
Metrics of the genome assemblies of *Alces alces* hap1 and hap2. The BlobToolKit Snailplots show N50 metrics and BUSCO gene completeness. The two outermost bands of the circle signify GC versus AT composition at 0.1% intervals, with mean, maximum, and minimum. The third outermost shows the N90 scaffold length, while the fourth is N50 scaffold length. The line from middle to second outermost band shows the size of the largest scaffold. All the scaffolds are arranged in a clockwise manner from largest to smallest, and shown in darker gray with white lines at different orders of magnitude, shown as a scale on the radius. The light gray shows the cumulative scaffold count. The scale inset in the lower left corner shows the total amount of sequence in the whole circle, and the fraction of the circle encompassed in the largest scaffold.

When compared to a k-mer database of the Hi-C reads, hap1 had a k-mer completeness of 96.2%, hap2 of 92.2%, and combined they have a completeness of 98.1%. Further, hap1 had an assembly consensus quality value (QV) of 49.5 and hap2 of 49.9, where a QV of 40 corresponds to one error every 10,000 bp, or 99.99% accuracy compared to a k-mer database of the Hi-C reads (QV 69.7 and 69.6, respectively, compared to a k-mer database of the HiFi reads) (Table 2). A total of 21,496 and 20,498 protein-coding genes were annotated in hap1 and hap2, respectively (Table 2).

Plotting the synteny of the assemblies generated here and other available assemblies of European moose (GCA_964270355.1, Elg.assembly.v.1.0; GCA_015832495.2, NRM_Aalces; and GCA_007570765.1, GSC_moose), visualises the different sequencing approaches used (Figure 3). These assemblies also have varying lengths in scaffolds and contigs (Table 3).

**Table 3:**
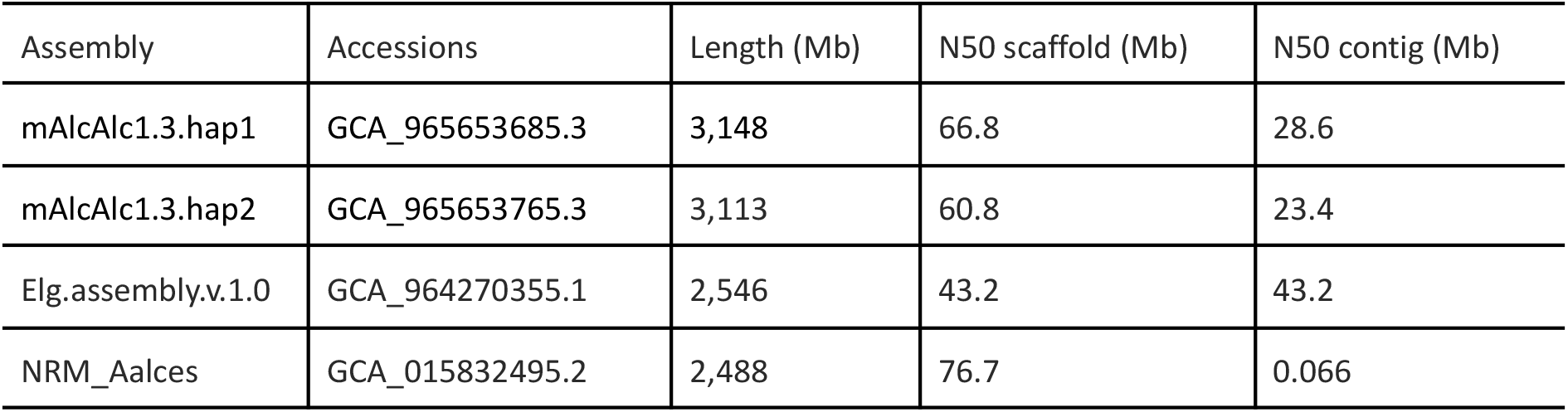
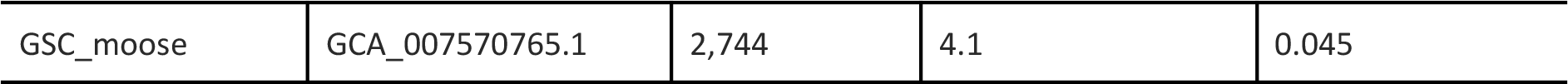
Genome statistics for available European moose genome assemblies.

| Assembly | Accessions | Length (Mb) | N50 scaffold (Mb) | N50 contig (Mb) |
| --- | --- | --- | --- | --- |
| mAlcAlc1.3.hap1 | GCA_965653685.3 | 3,148 | 66.8 | 28.6 |
| mAlcAlc1.3.hap2 | GCA_965653765.3 | 3,113 | 60.8 | 23.4 |
| Elg.assembly.v.1.0 | GCA_964270355.1 | 2,546 | 43.2 | 43.2 |
| NRM_Aalces | GCA_015832495.2 | 2,488 | 76.7 | 0.066 |
| GSC_moose | GCA_007570765.1 | 2,744 | 4.1 | 0.045 |

**Figure 3:**
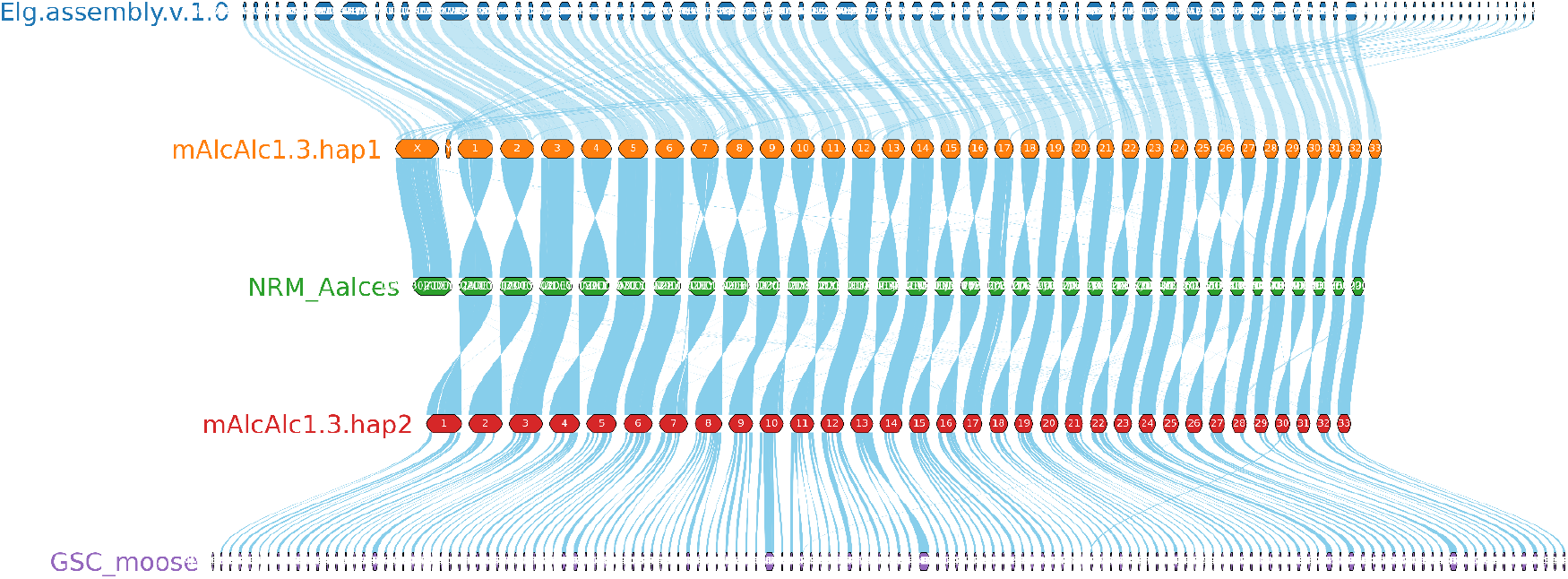
Synteny plot of available European moose genome assemblies. The three assemblies scaffolded with Hi-C data (NRM_Aalces and the two pseudo-haplotypes generated here [mAlcAlc1.3.hap1 and mAlcAlc1.3.hap2]) display chromosome-length scaffolds, while the two other assemblies (Elg.assembly.v.1.0 and GSC_moose) are fragmented. The new European genome assembly has 68 chromosomes, as expected.

## Discussion

The genome assembly presented here represents a chromosome level and pseudo-haplotype-resolved assembly more complete than any moose genome assemblies generated so far (Table 3). In comparison, the first *de novo* reference genome assembly for European moose, NRM_Aalces, generated using Illumina short-read sequencing, had a much shorter contig N50 at 0.066 Mb and a scaffold N50 of 1.7 Mb (Dussex et al., 2020). Dussex et al. (2023) later improved the original NRM_Aalces assembly by adding Hi-C scaffolding, achieving a chromosome-scale scaffold N50 of 76.7 Mb (Table 3). However, it is not indexed as a chromosome-level assembly in INSDC (International Nucleotide Sequence Database Collaboration), and it is missing chromosome Y since it was generated from a female (Figure 3). However, Dussex et al. (2023) does mention that they identified a Y chromosome, but that particular sequence (HiC_scaffold_21) aligns well with chromosome 19 in our assemblies, with chromosome Y identified separately. The GSC_moose assembly was sequenced with only Illumina short reads, and is substantially more fragmented than the other assemblies. The Elg.assembly.v.1.0 was assembled from Oxford Nanopore data without any long range information such as Hi-C, and does in some cases recover complete chromosomes, but a majority is fragmented into multiple sequences (Figure 3), although it has longer contigs than hap1 and hap2 (Table 3).

The moose is a heavily harvested species of major impact to rural economies (Storaas et al., 2001). Being a heat-sensitive species, recruitment is declining rather fast in the southern distribution ranges (Holmes et al., 2021; Lenarz et al., 2009; Murray et al., 2006), which already limit harvest rates and income to these economies. We need more insight into how rapid moose can adapt to the current warming, and whether the heavy and selective harvesting of moose is causing evolutionary changes. Changes in allele frequencies in moose were suggested to be attributed to selective harvesting (Dussex et al., 2023). However, teasing apart the relative role of climate warming and selective harvest for moose evolution is challenging, and these drivers may potentially cause divergent selection limiting adaptive capacity of moose. Therefore, this new genome assembly will be an important resource for accurate variant calling and for detecting structural variation in moose populations. Structural variation in moose will have impacts for choice of reference genome to use in further research. Moose in Europe and most parts of Asia have a 68-chromosome set, while North America and the easternmost parts of Asia have a 70-chromosome set. Hence, our reference genome provides an excellent starting point for studies into potential adaptive effects of warming, selective harvesting and possibly other drivers mainly in Europe and western parts of Asia.

## Acknowledgments

This project received data management and infrastructure support from ELIXIR Norway, supported by the Research Council of Norway’s grant 270068, the University of Bergen, the University of Oslo, the Arctic University of Norway in Tromsø, the Norwegian University of Science and Technology, and the Norwegian University of Life Sciences: NMBU. The authors acknowledge support from the National Infrastructure for High Performance Computing and resources provided by Sigma2 as well as Data Storage in Norway (project NN8013K) for computational work. The Norwegian Sequencing Centre generated the sequencing data used in this project (http://sequencing.uio.no). The authors would like to thank the Kraakstad hunting team and Kristian Sæther for collecting the sample.

## Data availability

Data generated for this study are available under ENA BioProject PRJEB90379. Raw PacBio sequencing data for the European moose (ENA BioSample: SAMEA117970373) are deposited in ENA under ERR15004688, whereas Illumina Hi-C sequencing data are deposited in ENA under ERR14902783. Pseudo-haplotype one can be found in ENA at PRJEB88053, whereas hap2 is PRJEB90378.

The gene annotations (and script for plotting synteny) are available at https://doi.org/10.5281/zenodo.20269287.

