## Supplementary information for "A chromosome-level reference genome of the largest cervid species - the European moose (*Alces alces*; Linnaeus, 1758)"

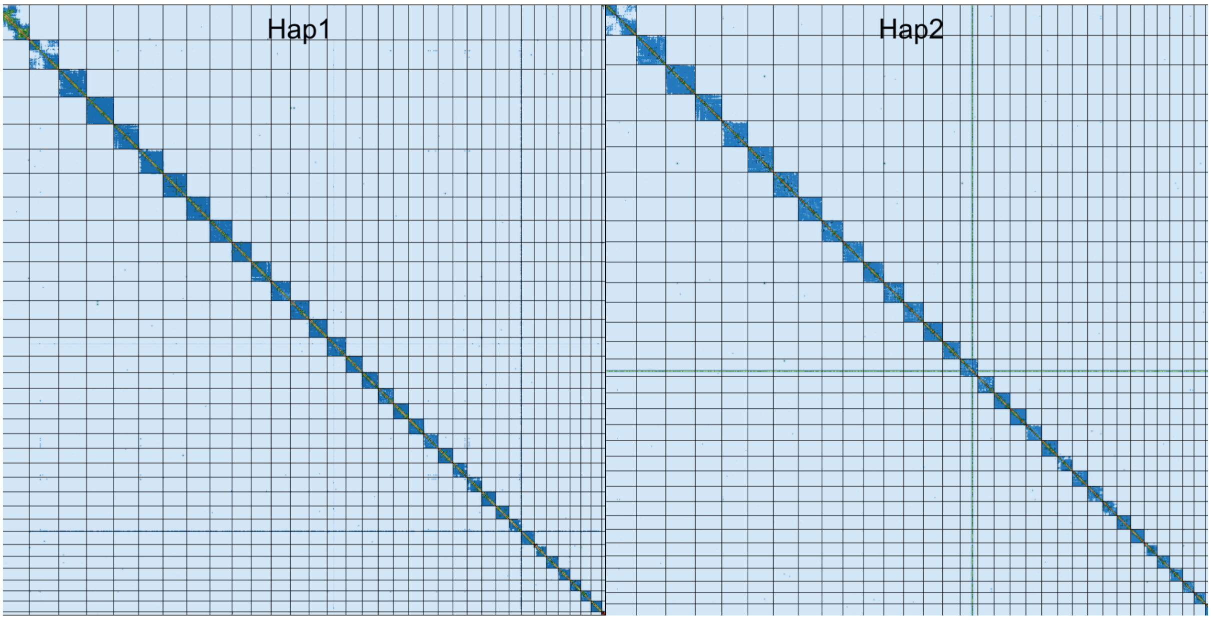


**Supplementary Figure 1: Hi-C contact maps for *Alces alces* hap1 and hap2.** The contact map displays interaction frequencies between genomic regions, where darker shades represent a higher number of Hi-C contacts. The axes correspond to the coordinates along each assembly. The Hi-C contact maps were generated by mapping the Hi-C reads to the haplotype genomes using BWA-mem, generating a contact map using PretextMap and visualized using PretextSnapshot.

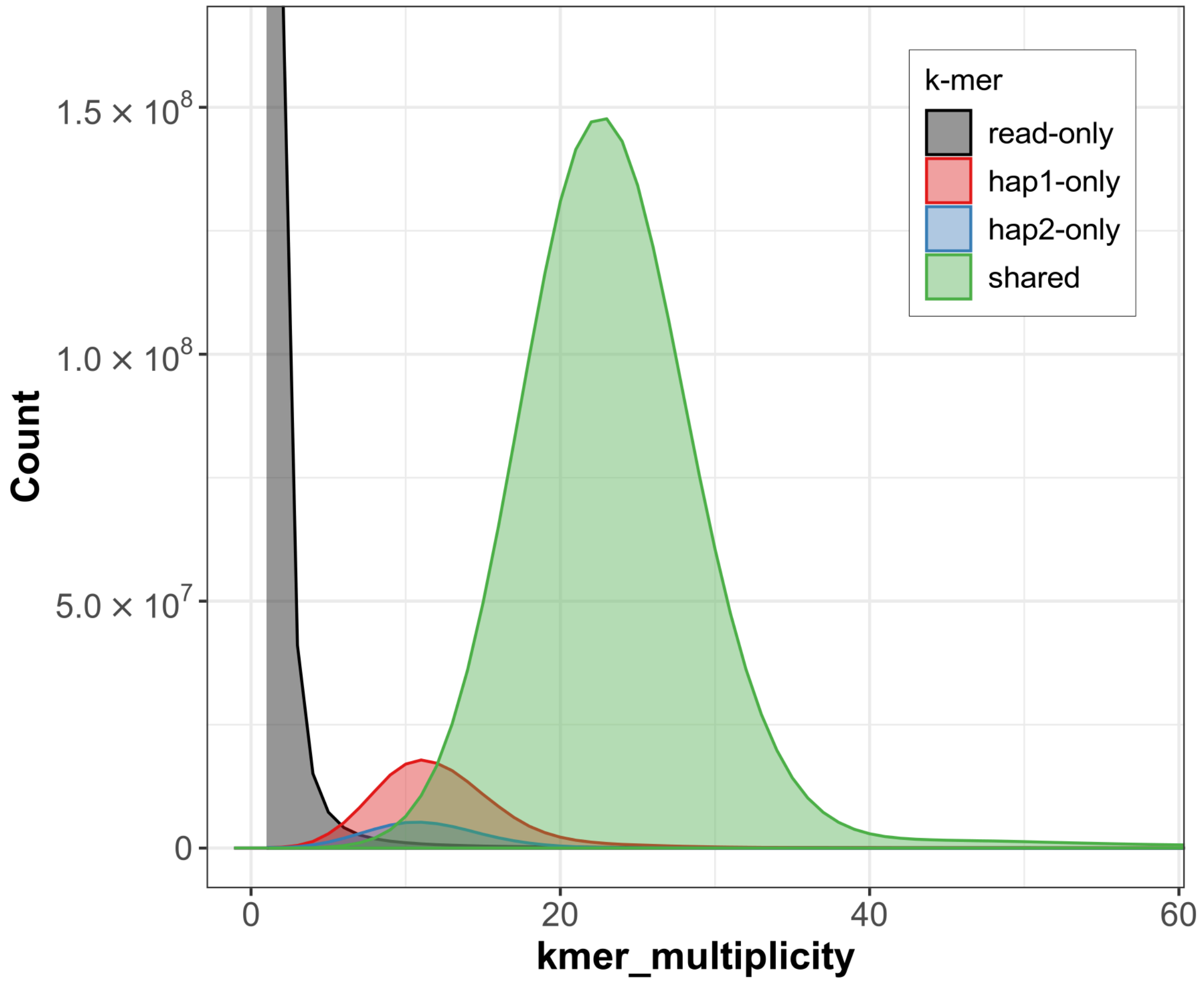


**Supplementary Figure 2: *K*-mer spectra of HiFi reads for *Alces alces*.** Distributions of *k*-mers found only in the reads (black), only in haplotype 1 (red), only in haplotype 2 (blue), or in both haplotypes (green). The x-axis represents the number of unique *k*-mers, while the y-axis represents the *k*-mer multiplicity (how often the *k*-mer is found in the set of HiFi reads).

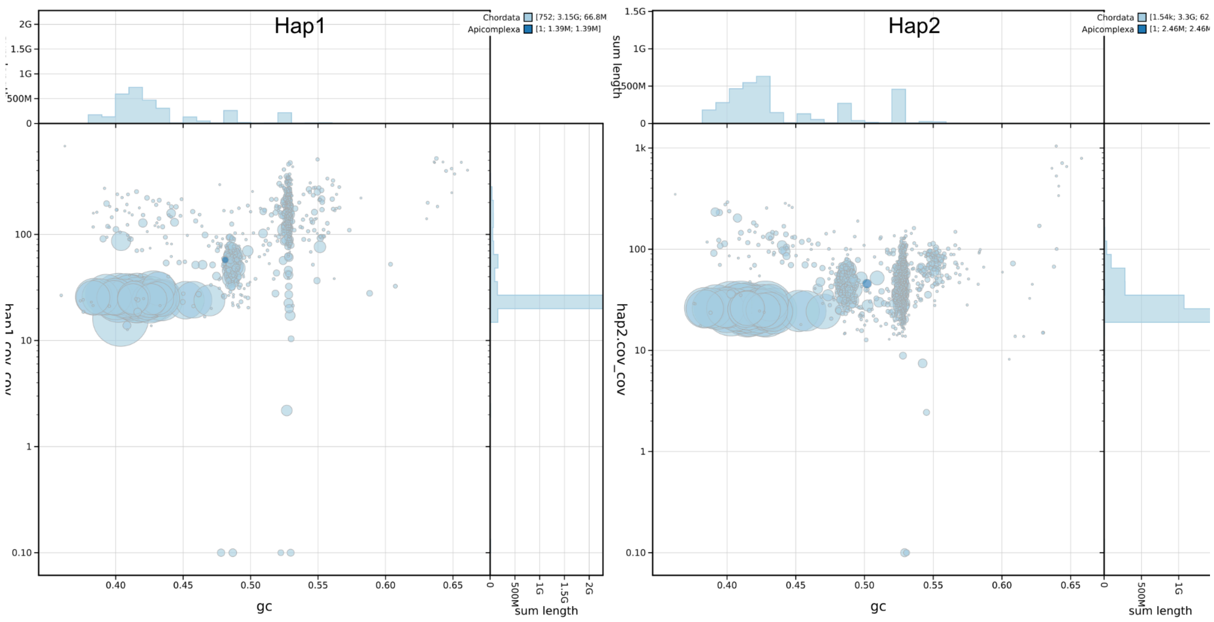

**Supplementary Figure 3: Coverage vs GC plots of *Alces alces* hap1 and hap2.** The BlobToolKit Blobplots depicts each scaffold as a dot based on the GC content (%GC, x-axis) and coverage (Y-axis). Size of the dots correspond to scaffold length. Dots are colored based on assigned taxonomy. Histograms of sequence lengths within a certain %GC range or coverage range are depicted on the top and right respectively.
